# Loss of cell-identity DNA methylation tracks division history, not DNA damage

**DOI:** 10.64898/2026.11.03.763914

**Authors:** Naor Sagy, Nadav Benedek, Daniel Z Bar

**Affiliations:** Department of Oral Biology, Gray Faculty of Medical & Health Sciences, The Goldschleger School of Dental Medicine, Tel Aviv University, Tel-Aviv, Israel; Reichman University, Israel

## Abstract

One influential proposal holds that DNA damage drives aging by degrading the epigenome, so that a cell loses the information that specifies what it is. We test that specific proposed driver, not the broader idea that aging involves a loss of epigenetic information. Cell-type-unique CpGs are a direct place to test it. They are the sites where a cell’s methylome differs most from every other cell type, so they are where its identity information can be accurately measured. We built such a panel for human skin fibroblasts against twenty primary cell types, and found that most of its 9,150 CpGs behave as a record of proliferation rather than identity. Their methylation state in fibroblasts is shared with continuously proliferating cell lines, consistent with the bulk hypomethylation of late-replicating chromatin that accumulates in culture, and this layer must be separated from identity marks before identity loss can be tested. We then expressed every perturbation as the fraction of the panel’s identity information lost beyond baseline-matched background CpGs, using a Bernoulli cross-entropy between the fibroblast code and a generic comparator cell. In 24 matched fibroblast cultures, replicative exhaustion cost 24.3% of the identity information (95% CI 20.2 to 28.7) and RAS activation cost 5.4% (4.6 to 6.1), both at p = 1.2 × 10⁻⁷. A senescence-inducing dose of X-rays cost nothing: −0.5% (−0.7 to −0.2), with identity CpGs if anything moving slightly further from other cell types than matched background did. The design could detect a loss of 0.4%, about one sixtieth of the exhaustion effect, so the X-ray result is a bounded negative rather than an absence of power. Doxorubicin-induced and irradiation-induced senescence did not move the panel either. Clones that kept dividing after irradiation moved further from other cell types rather than collapsing toward them. In human fibroblasts, loss of identity information tracks prolonged division, not acute DNA damage.

**In brief:** We directly test the proposal that DNA damage erodes epigenetic cell identity. In the same human fibroblast cultures, replicative exhaustion costs a quarter of the identity information carried by fibroblast-unique CpGs. A senescence-inducing dose of X-rays costs none beyond what baseline-matched background CpGs show, in a design that would have detected a loss sixty times smaller than the exhaustion effect. Identity loss tracks prolonged division, not the acute damage dose.

**Highlights:**

- Most CpGs unique to a cultured cell type record division history rather than lineage identity, and must be separated before identity loss can be measured
- A cross-entropy score, corrected for baseline-matched background, expresses each perturbation as the fraction of a cell’s identity information that was lost
- Replicative exhaustion costs 24% of the identity information at fibroblast-unique CpGs; a senescence-inducing X-ray dose costs none, with a detection limit of 0.4%
- Oncogene activation erodes identity at these sites, damage that arrests the cell does not, and damage followed by continued division moves them further away from other cell types

## Introduction

DNA damage has a strong claim to being a primary driver of aging. Somatic genomes accumulate lesions throughout life, and the machinery that repairs them declines in capacity. Humans with inherited defects in that machinery age early and segmentally: Werner syndrome, Cockayne syndrome, trichothiodystrophy and XFE progeroid syndrome each compress a recognisable subset of aging phenotypes into a short lifespan (Hoeijmakers 2009). Mice engineered with the same lesions do the same. *Ercc1* hypomorphs and *Xpd* mutants shorten lifespan and reproduce much of the aged transcriptional and physiological phenotype (Niedernhofer et al. 2006; de Boer et al. 2002). Genome instability is listed first among the hallmarks of aging for these reasons (López-Otín et al. 2013; López-Otín et al. 2023). Damage also drives cellular senescence, which accumulates with age and spreads dysfunction through the senescence-associated secretory phenotype (d’Adda di Fagagna 2008; Di Micco et al. 2021).

A second, more specific claim has been built on top of the first. The proposal is that damage matters not mainly because of the mutations it leaves behind, but because repairing it pulls chromatin regulators away from their normal sites and leaves the epigenome permanently altered. Early support came from the observation that double-strand breaks redistribute SIRT1 away from its target promoters, producing an aging-like expression profile (Oberdoerffer et al. 2008). The strongest version of the argument is the ICE mouse, in which repeated non-mutagenic double-strand breaks accelerate epigenetic clocks and phenotypic aging, and partial reprogramming reverses part of it (Yang et al. 2023). The authors frame this process as a loss of epigenetic information, in which a cell gradually loses the marks that specify its lineage. The model is compelling because it offers a concrete mechanism that is, in principle, reversible.

Testing this hypothesis requires a readout of cell identity rather than chronological age. Epigenetic clocks are accurate age predictors, but they were not designed to measure lineage identity and can advance for reasons unrelated to a cell losing its type (Horvath 2013). Tissue-unique and cell-type-unique CpGs provide a direct readout. Defined as the loci where one cell type sits clearly outside the methylation range of all others, they carry the specific information that distinguishes that lineage from the rest. If DNA damage erodes epigenetic identity, these loci should drift toward the methylation values held by other cell types. We previously built an atlas of such sites across 22 human tissues to test questions of this kind (Sagy et al. 2025).

Our own published work hints that damage might indeed be the driver. In liver, the collapse of tissue-unique CpGs toward the population mean scales continuously with histological fibrosis stage across its full clinical range (Spearman ρ = 0.70, n = 341; (Sagy et al. 2025; Shtark et al. 2026)), and the same sites regress with age in healthy mouse liver (87.4% of sites). In colon, tissue-unique sites track chronological age closely enough to power a tissue-specific clock, and that clock runs further ahead in colitis and in colonic polyps (Sagy et al. 2026). Fibrosis, chronic inflammation and polyp formation are all states of elevated genotoxic burden. Read on their own, these results are consistent with cumulative DNA damage eroding identity marks, and that reading is what motivated the present study.

Yet bulk tissue data alone cannot isolate the mechanism. In a tissue, damage never travels alone. Injury brings inflammation, compensatory proliferation, changes in cell composition and clonal selection, and each of these can move a bulk methylation panel without any cell having lost its identity program. Nothing in an observational cohort separates them. A causal answer needs a system where the dose is set by the experimenter, the cell type is pure, and division can be varied. Primary human fibroblasts are such a system. They can be irradiated at a defined dose, pushed into senescence by distinct triggers, passaged to exhaustion, derived as single-cell clones and followed for a known number of population doublings, all within a single, pure cell type.

Here we test the damage hypothesis in human fibroblasts, extensively and in one consistent framework. We first rebuilt a fibroblast-unique CpG panel against twenty primary comparator cell types and found that most of it records proliferation rather than identity, reflecting the bulk hypomethylation of late-replicating chromatin that marks any heavily divided cell, and must be separated out before identity loss can be measured. We then put every perturbation on a single normalized scale by asking what fraction of the panel’s identity information it costs beyond what baseline-matched background CpGs show, using a Bernoulli cross-entropy between the panel’s own cell type and a generic comparator cell. Across five public datasets we scored replicative exhaustion, oncogene-induced senescence, acute X-irradiation, doxorubicin-induced and irradiation-induced senescence, chronic low-dose irradiation, progeria, and single-cell-derived clones that kept dividing for about twenty doublings after a 2 Gy dose. We also scored one in vivo contrast, sun-exposed against sun-protected skin from the same people, in which keratinocyte identity does collapse; but a single cohort cannot separate ultraviolet damage from the inflammation, hyperproliferation and clonal replacement that accompany it, so we report that arm as supplementary material only and draw no conclusion from it (Supplementary Figures S4 and S5). The answer throughout is the same. Replicative exhaustion and oncogene activation cost identity information. Acute DNA damage costs none that we can detect, and damage followed by continued division moves the panel the other way.

## Results

### Measuring epigenetic identity loss at fibroblast-unique CpGs

A cell type’s identity is written most legibly at the CpGs where its methylome differs from every other cell type. We defined a CpG as fibroblast-unique if the mean of five primary skin fibroblast lines sat at least 0.1 beta units outside the 5th to 95th percentile range of twenty primary non-fibroblast cell types, using one value per cell type (Figure 1A; Methods; (Sagy et al. 2025)). This produced 9,150 CpGs, 6,621 uniquely low (UL) and 2,529 uniquely high (UH).

**Figure 1:**
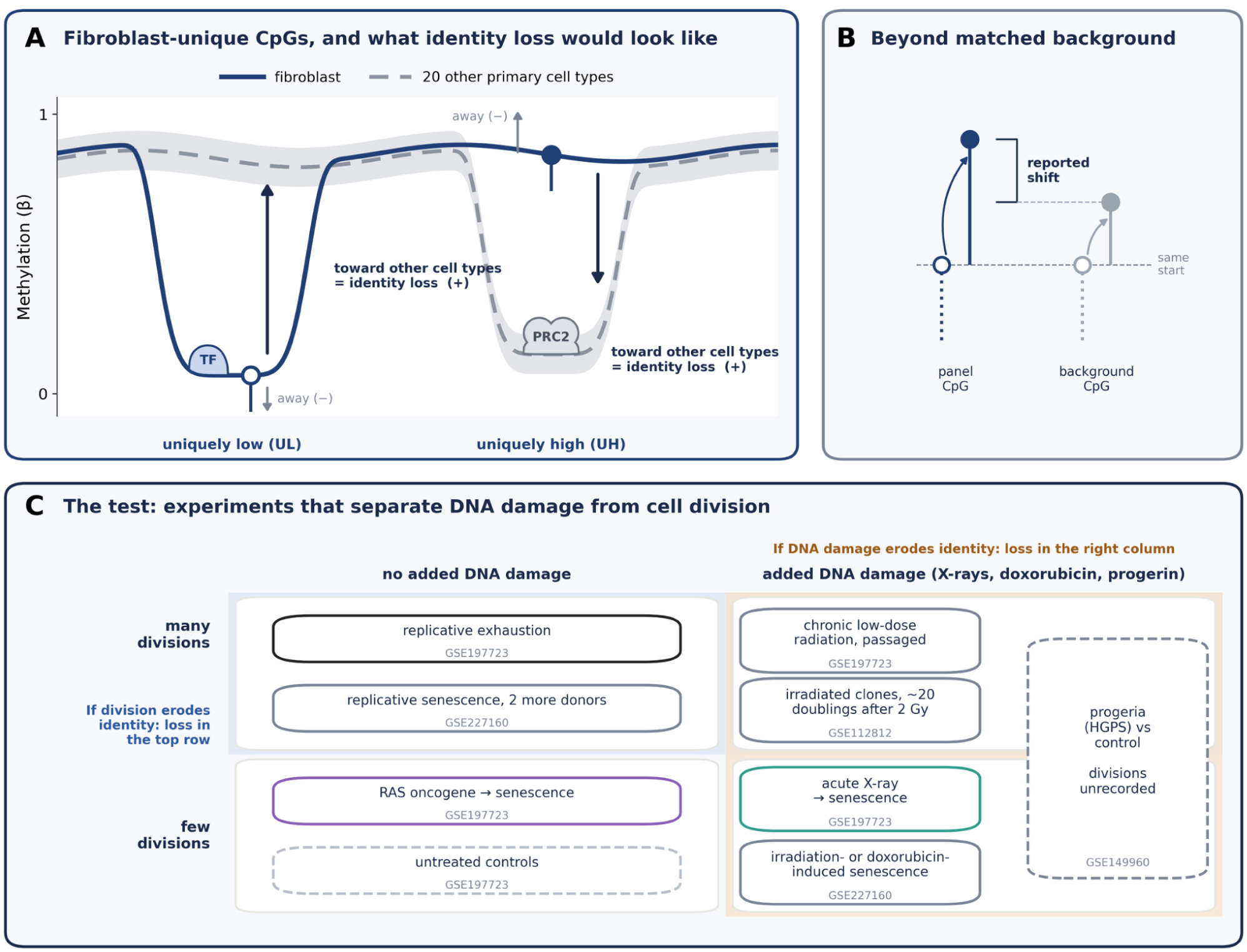
Measuring identity loss at fibroblast-unique CpGs, and the experiments that separate DNA damage from division. **(A)** Schematic of methylation along a stretch of genome in fibroblasts (solid) and in the 20 other primary cell types (dashed line, mean; band, range). A fibroblast-unique CpG is uniquely low (UL, open circle, unmethylated) or uniquely high (UH, filled circle, methylated): the fibroblast mean lies at least 0.1 beyond the 5th or 95th percentile of the other types (real examples in Figure 2B). TF, a transcription factor bound beside the unmethylated UL site; PRC2, Polycomb repressive complex 2, bound at the UH site in the other cell types, where it is unmethylated. Identity loss is movement toward the values the other cell types hold, upward at UL and downward at UH; movement away is not identity loss. In every figure positive means toward the other cell types. **(B)** Every shift is taken beyond 20 background CpGs per panel CpG that lie outside the CpGs being scored but start at the same methylation, because CpGs at extreme methylation drift toward the middle on their own. **(C)** The designs used, sorted by whether exogenous DNA damage was added and whether the cells went on to divide many times. GSE197723 splits one culture into an untreated control and replicative-exhaustion, RAS and acute X-ray arms (coloured as in Figure 2), 24 times. RAS acts through oncogene-induced hyperproliferation and replication stress before arrest, which itself triggers a DNA-damage response; “no added DNA damage” means no exogenous genotoxin. The progeria cultures and their controls were at early passage, but their population doublings are not recorded. If DNA damage drives identity loss, the loss should appear in the right column; if division does, in the top row.

If a fibroblast loses its identity, these CpGs should move toward the values the other cell types hold, upward at UL sites and downward at UH sites (Figure 1A). Movement the other way, away from the other cell types, is not identity loss. One correction is needed before either can be read. Panel CpGs sit at extreme beta by construction, and CpGs at extreme beta drift back toward the middle whether or not they carry identity. We therefore matched every panel CpG to 20 CpGs outside the scored set that start at the same beta, and every score in this paper is the panel’s shift beyond that background (Figure 1B; Methods). Throughout, positive means toward the other cell types.

Which perturbation moves these CpGs is only interpretable if DNA damage and cell division are varied separately (Figure 1C). The core design of (Kabacik et al. 2022), GSE197723, splits each of 24 primary fibroblast cultures four ways: an untreated control, a culture passaged to replicative exhaustion, a culture given a senescence-inducing dose of X-rays, and a culture carrying activated RAS (Kabacik et al. 2022). Scoring each culture against its own control removes donor, batch and chip. Exhaustion adds many divisions over months of culture and no exogenous damage, while X-rays add acute damage and stop division within about two weeks. Further datasets add replicative and damage-induced senescence in two more donors, chronic low-dose irradiation during passaging, progeria, and irradiated single-cell clones that kept dividing. If damage erodes identity, the loss should grow along the damage axis; if division does, it should grow along the division axis.

### Most fibroblast-unique CpGs record division, not identity

Scored as a single unsplit panel, these CpGs move in the opposite direction to what identity loss predicts. Under replicative exhaustion they moved away from the other cell types, by 0.154 at UL and 0.113 at UH, in all 24 cultures (Figure 2A). Read literally, an exhausted fibroblast would be a more distinctive fibroblast than a young one. RAS moved the panel the same way at about a quarter of the size. X-rays moved it toward the other cell types, consistently (24 and 21 of 24 cultures) but only slightly (+0.005 at UL, +0.002 at UH), about a thirtieth of the exhaustion shift.

**Figure 2:**
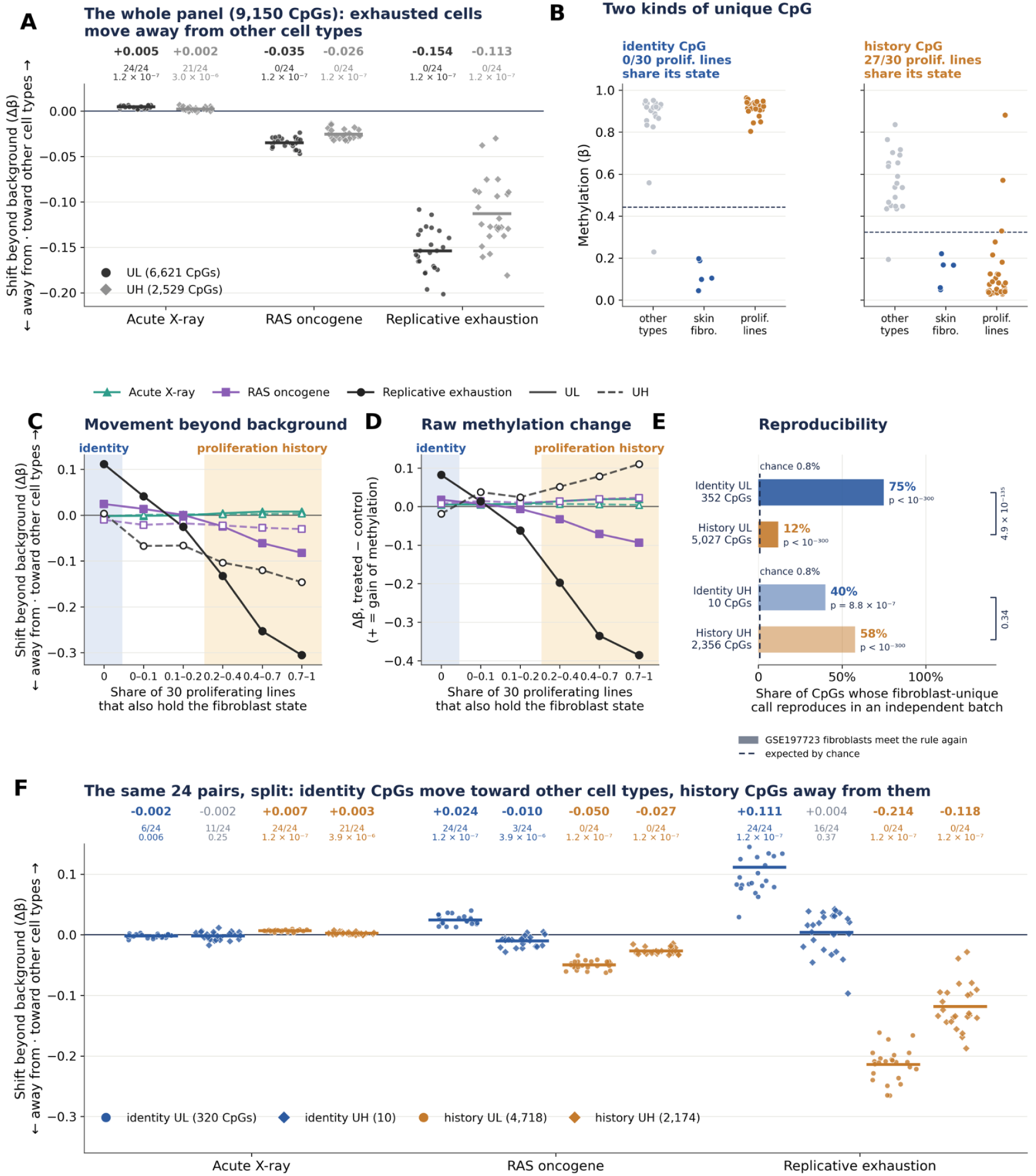
Most fibroblast-unique CpGs record division history rather than identity. Shifts are beyond baseline-matched background, in beta, positive toward the other cell types. **(A)** The whole panel (6,621 UL and 2,529 UH CpGs) in the 24 matched GSE197723 quadruplets, one point per culture paired with its own control. Above each group: mean shift, the number of the 24 pairs that moved toward the other cell types, and the Wilcoxon signed-rank p (1.2 × 10⁻⁷ is the smallest p possible with 24 pairs). Exhaustion and RAS move the whole panel away from the other cell types. **(B)** An identity CpG and a proliferation-history CpG in GSE40699: the 20 other primary cell types, the five skin fibroblast lines, and 30 continuously proliferating lines. At the history CpG most proliferating lines share the fibroblast state. **(C)** Shift beyond background against the share of the 30 proliferating lines that also hold the fibroblast state (bins of panel CpGs; UL solid, UH dashed). Under exhaustion, CpGs no proliferating line shares (identity, blue band) move toward the other cell types (+0.111 at UL), and CpGs most of them share (amber band) move away (−0.306), with a monotonic gradient between. RAS follows the same gradient at smaller size; acute X-ray is nearly flat. **(D)** The same bins in raw methylation change: the identity end gains methylation at UL, and the proliferation-history end loses large amounts of it, in the pattern of partially methylated domains. **(E)** Reproducibility: the share of each layer’s CpGs at which fibroblasts from an independent batch (GSE197723) again meet the fibroblast-unique rule against the 20 ENCODE comparators. The dashed line marks the share expected by chance (the same test at every array CpG outside the panel, 0.8%), with the binomial p against it beside each bar; brackets give Fisher’s exact p for identity against history CpGs of the same direction. **(F)** The quadruplets of panel A split into identity (blue) and proliferation-history (amber) CpGs, UL as circles and UH as diamonds; numbers as in panel A.

The reason is that a cultured cell’s methylome carries a second signal at the same kind of site. Late-replicating, lamina-associated chromatin loses methylation in bulk as cells divide, forming partially methylated domains, while Polycomb-target CpG islands gain it, and both scale with the number of divisions rather than with lineage (Zhou et al. 2018). A passaged fibroblast is a genuine outlier at these sites, so they enter a uniqueness panel by construction without encoding fibroblast identity.

We separated the two with the simplest proxy available in the same reference series. For each panel CpG we asked how many of the 30 continuously proliferating cell lines in the reference series, the most heavily divided cells in the dataset, also sit beyond the fibroblast threshold (Figure 2B). CpGs that none of them share we call identity CpGs (352 UL, 10 UH), and CpGs that at least a fifth of them share we call proliferation-history CpGs (5,027 UL, 2,356 UH); the remainder belong to neither layer. These are operational labels: sharing a state with heavily divided lines marks a CpG as a plausible record of proliferation, but we did not test these loci directly for membership of partially methylated domains, lamina-associated domains or late-replicating chromatin. The split is sharp in every way we can check it. Under passaging, the shift changes monotonically with the share of proliferating lines, from toward the other cell types at the identity end to strongly away from them at the history end (Figure 2C). The raw changes behind it differ in kind: the identity layer gains methylation at UL, while the history layer loses large amounts of it (Figure 2D). And the layers differ in reproducibility (Figure 2E). In an independent batch (GSE197723), fibroblasts again meet the fibroblast-unique rule against the twenty ENCODE comparators at 75% of identity UL CpGs but at only 12% of proliferation-history UL CpGs (p = 5 × 10⁻¹³⁵). A CpG chosen at random meets it 0.8% of the time, so both layers are genuine features of cultured fibroblasts; what separates them is that the identity CpGs reproduce six times as often. At UH the layers do not differ (40% and 58%, p = 0.34), though the identity layer has only 10 UH CpGs.

Split this way, the same 24 cultures tell a coherent story (Figure 2F). Under exhaustion the identity CpGs moved toward the other cell types (+0.111 at UL, all 24 cultures, p = 1.2 × 10⁻⁷), while the history CpGs moved away from them (−0.214 at UL, −0.118 at UH). RAS repeated the pattern at UL on a smaller scale (identity +0.024, history −0.050). Under X-rays the identity layer moved slightly away from the other cell types (−0.002 at UL, p = 0.006), so the small whole-panel shift came from the history layer. Weighted by CpG count, the two layers moved in opposite directions under exhaustion, the identity layer 0.108 toward the other cell types and the history layer 0.184 away from them, which is why the unsplit panel reads as net divergence.

Nothing about this result depends on using the proliferating lines to split the panel after construction rather than excluding them at the start. Building the panel with the same 30 lines among the comparators instead, so that proliferation CpGs never enter it, yields 1,154 UL CpGs that reproduce the identity-layer result (replicative exhaustion +0.082, RAS +0.018, X-ray −0.002) and include 348 of the identity CpGs, at the cost of leaving only 25 UH CpGs (Methods). We kept the layered panel because it preserves both directions. We also tried three rules that use no proliferating-line data at all, selecting on how far outside the comparator range a CpG sits, on how tightly the comparators themselves cluster, or on whether the unique call reproduces in an independent batch. None of them isolates the identity layer, and under all three the panel still moves away from the other cell types under passaging (Methods). Everything below is reported for the identity layer, with the proliferation-history layer alongside it as a control.

### Division costs identity information, and acute damage costs none

A shift in beta is hard to compare across panels of different size and different baseline methylation, and it needs a separate sign convention for UL and UH. We therefore converted every methylation shift into the quantity the hypothesis directly concerns: the fraction of cell-identity information lost (Figure 3A). Treating each CpG as a Bernoulli variable, we described each sample under two codes, the panel’s own cell type and a generic other cell taken as the mean of the comparator primary types. The extra code length of the identity model is linear in beta, so any contrast reduces to a weighted sum of methylation differences, and dividing by the cost of moving every CpG the whole way to the generic cell puts it on a common scale (Methods). Zero means unchanged, one means every CpG moved the whole way to the generic cell’s value, and negative means moved further away. Equivalently, it is the average over CpGs of how far each moved from the fibroblast value toward the generic value, each CpG weighted by how strongly it separates the two (Figure 3A). It is therefore an information-weighted displacement toward the generic cell at these CpGs, not a measure of everything that makes a cell a fibroblast. Because the weight carries the sign, UL and UH become two halves of one number, each weighted by the information it carries; for the identity layer that is 98% UL, and Supplementary Figure S3 shows the two halves separately for every contrast. As before, movement only counts beyond the matched background: from each contrast we subtract the loss the panel would have shown had every CpG moved like its baseline-matched background CpGs (Methods).

**Figure 3:**
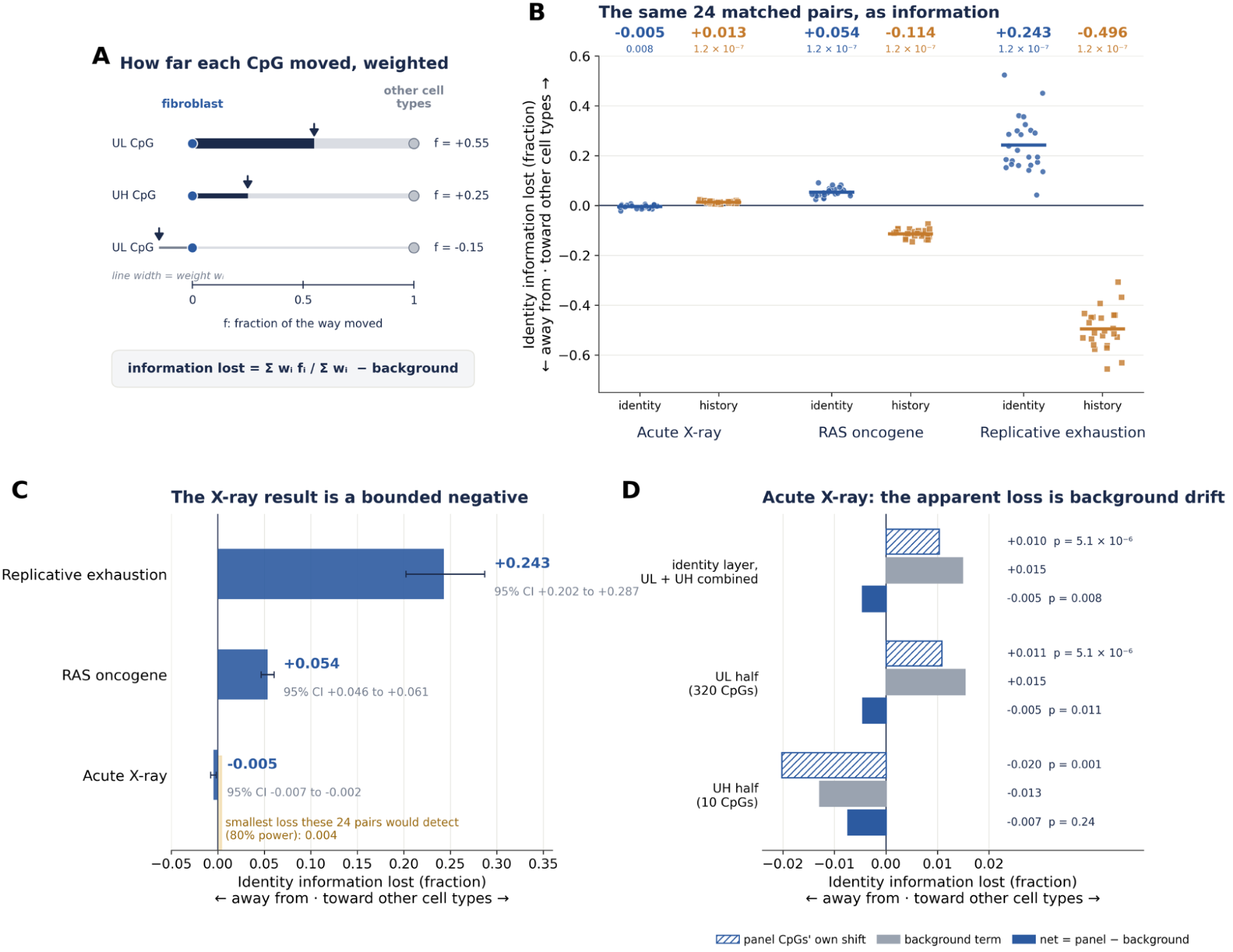
Division costs identity information; acute DNA damage costs none. **(A)** What the fraction measures. For each CpG, f is the fraction of the way a sample moved from the fibroblast value toward the mean value of the other cell types; f = 0 is unchanged, f = 1 is the other cell types’ value, and negative f is further from them. UL and UH CpGs sit on the same scale. The identity information lost is the average of f over CpGs, each weighted by its Jeffreys term w (line width), which is largest where fibroblasts and the other cell types differ most, minus the same quantity computed from each CpG’s 20 matched background CpGs (Methods). **(B)** The 24 GSE197723 quadruplets, one point per culture: identity information lost in the identity layer (blue) and in the proliferation-history layer (amber). Above each group: mean and Wilcoxon signed-rank p. **(C)** The identity layer with 95% bootstrap intervals. The shaded band marks the smallest loss the 24 pairs would detect with 80% power (0.004). **(D)** Acute X-ray, identity layer: the panel CpGs’ own shift, the background term (each panel CpG’s 20 background CpGs, weighted as that CpG is), and their difference, for UL and UH combined and for each half. Without the subtraction the UL and UH halves move in opposite directions, because both gain methylation; net of background, neither moves toward the other cell types.

The panel is where the identity information is concentrated. The 9,150 panel CpGs are 1.9% of the array but carry 17.7% of its total identity information, with a mean absolute weight of 1.72 against 0.41 for all other CpGs. No small subset drives the total: the participation ratio of the weights is 5,734 of 9,150. Clipping the reference probabilities at 0.01 instead of 0.02 changes any contrast by at most 0.0001. We note what this measure is not. Within a panel and direction it correlates with the shift beyond background at Spearman 0.97 to 1.00 across contrasts, so it is a principled reweighting of the same measurement rather than independent evidence. What it provides is a single signed, normalized value per contrast, allowing perturbations of very different kinds to be compared on one axis.

On this scale, replicative exhaustion cost 24.3% of the fibroblast identity information (95% CI 20.2 to 28.7, p = 1.2 × 10⁻⁷, 24 pairs) and RAS activation cost 5.4% (4.6 to 6.1, p = 1.2 × 10⁻⁷). X-irradiation cost nothing: −0.5% (−0.7 to −0.2, p = 0.008) (Figure 3B). All 24 cultures lost identity information under exhaustion and under RAS, while under X-rays 18 of 24 moved slightly the other way. The proliferation-history layer moved the other way on the same scale, by 0.496 under exhaustion and 0.114 under RAS. In the same cells, the identity information lost tracks prolonged division and oncogene activation, not the acute DNA-damage dose.

A negative result is only informative if the design could have detected the positive, and two features of these data make the bound explicit (Figure 3C). First, the X-ray estimate is tight. Its 95% interval runs from −0.7% to −0.2%, so it excludes any loss on this score, and it amounts to −0.005 bits per CpG against 0.290 bits per CpG for replicative exhaustion. The small negative value is nominally significant (p = 0.008), but it is no larger than the design’s detection limit, and we do not read it as a gain of identity. Second, with 24 matched pairs and the observed per-pair variability, this design has 80% power to detect a loss of 0.004 on this score, an effect about one sixtieth the size of replicative exhaustion. The X-ray arm is therefore a bounded negative rather than an untested claim of no effect.

The background subtraction decides this arm (Figure 3D). Scored without it, X-rays would appear to cost 1.0% of the identity information (p = 5.1 × 10⁻⁶). But CpGs at the same baseline methylation drifted toward the generic cell slightly further than the identity CpGs did: the background term was +0.015, larger than the panel’s own unadjusted shift of +0.010. Without the subtraction the two halves of the identity layer also disagree. Under X-rays its UL and UH CpGs both gained about 0.005 in methylation, which reads as a loss at UL (+1.1%) and a gain at UH (−2.0%, p = 0.001). A uniform shift does that; a move toward another cell type would push both halves the same way. Under exhaustion and RAS the background term ran opposite to the panel (−0.050 and −0.014), so subtracting it enlarges both losses, and neither is regression to the mean.

### The pattern holds across triggers, donors and designs

Figure 4A places every fibroblast contrast on the same scale. Three further arms test the GSE197723 result from different directions.

**Figure 4:**
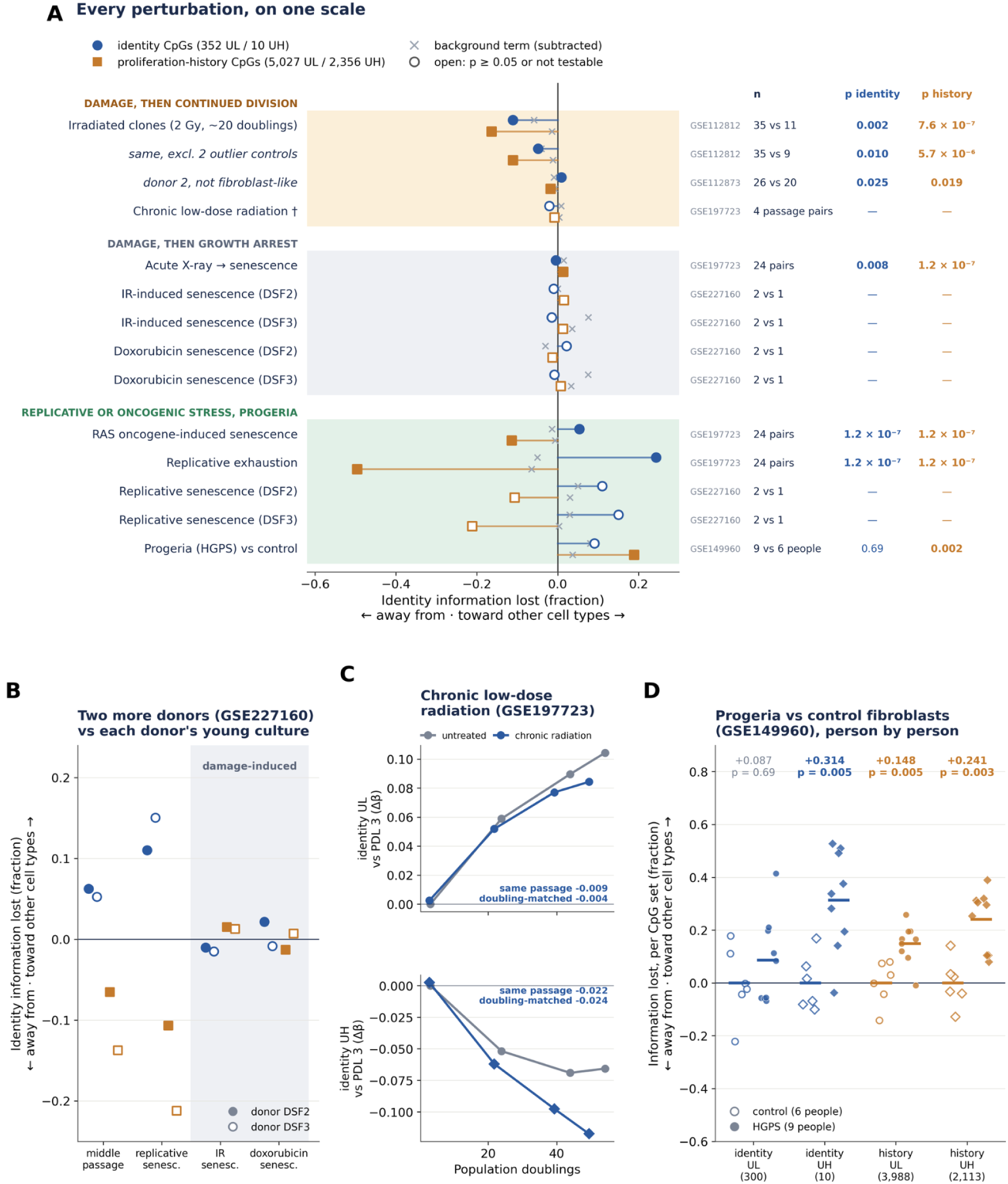
Every perturbation on one scale. **(A)** Identity information lost in every fibroblast contrast, for the identity layer (blue circles) and the proliferation-history layer (amber squares), with the subtracted background term (×). Filled markers are p < 0.05 at the sample level (Wilcoxon signed-rank for paired designs, Mann-Whitney otherwise); open markers are not significant or not testable. † The chronic-radiation cultures sit 5 to 10% lower in population doublings than their controls; panel C shows the doubling-matched comparison. **(B)** GSE227160: two further dermal fibroblast donors (filled DSF2, open DSF3), each against its own young culture. With one reference culture per donor these values are descriptive. **(C)** Chronic low-dose irradiation over passaging, identity layer: shift beyond background relative to the untreated culture at PDL 3, in beta. The numbers give the chronic-radiation shift against untreated cultures of the same passage, and after interpolating the untreated series to the irradiated cultures’ doublings. **(D)** Progeria (GSE149960), nine HGPS and six control people: information lost separately for the UL and UH halves of each layer, with the number of CpGs scored in brackets; mean and Mann-Whitney p. The 10 identity UH CpGs carry under 2% of the identity layer’s information.

Replicative senescence reproduces in independent donors; damage-induced senescence does not. GSE227160 contains two dermal fibroblast donors taken to replicative senescence, to doxorubicin-induced senescence and to irradiation-induced senescence, each with its own young culture as reference (Figure 4B). Replicative senescence cost 11.0% and 15.0% of the identity information in the two donors, less than the 24.3% of the matched-quadruplet experiment but in the same direction, and a middle-passage culture sat between the young reference and the senescent endpoint in both (6.3% and 5.2%). The two damage-induced triggers gave +2.2% (doxorubicin) and −1.0% (irradiation) in one donor and −0.8% and −1.5% in the other, with no consistent direction across donors or triggers. With one reference culture per donor these values are descriptive, and we did not test them.

Chronic low-dose irradiation across a passaging series barely moved the identity layer relative to untreated cultures of the same passage: −0.009 at UL and −0.022 at UH beyond background, both slightly away from the other cell types, or −2.1% of the identity information (Figure 4C). The irradiated cultures sit 5 to 10% lower in population doublings than their controls, and interpolating the untreated series to matched doublings gives −0.004 and −0.024. On the unsplit panel the same adjustment slightly enlarges a near-zero shift rather than removing one (Supplementary Figure S1C). Its four pairs are passage points in one series rather than independent experiments, so we report it as uninformative rather than negative. Acute DNA damage therefore costs no detectable identity information, while the available chronic-radiation series is too small to test chronic exposure on its own.

Hutchinson-Gilford progeria is the obvious test case for damage-driven identity loss, since progerin accumulation causes persistent DNA damage alongside its structural effects on the nuclear lamina. Comparing nine HGPS to six control dermal fibroblast cultures (two parental and four age-matched lines, all at early passage; Methods), the proliferation-history layer lost 18.9% of its information (95% CI 11.4 to 26.7, p = 0.002) and the identity layer lost 9.1%, which was not significant (−4.8 to 23.7, p = 0.69) (Figure 4A). Against the four age-matched controls alone, the history layer lost 21.0% (p = 0.006) and the identity layer 2.6% (p = 0.94). Both layers moved toward the other cell types, and the full panel’s shifts beyond matched background were among the largest in the study (+0.051 at UL, +0.091 at UH; Supplementary Figure S1E).

This is not evidence for the damage hypothesis. Within the identity layer, the 10 UH CpGs did move, losing 31% of their information (p = 0.005) as the history layer’s UH CpGs also did, but they carry under 2% of the layer’s information; the UL CpGs that carry the rest did not (8.7%, p = 0.69) (Figure 4D). At nine versus six individuals the arm has 80% power only for a loss above 0.228, close to replicative exhaustion’s own effect, so we read it as underpowered rather than negative. More informative is the pattern across layers. In every culture perturbation where identity collapses, the two layers dissociate: exhausted and RAS-arrested cultures move the history layer away from the other cell types while the identity layer moves toward them. In progeria both move toward the other cell types, and only the history layer’s loss is significant, although the difference between the two layers is not (p = 0.33). If anything, that points to the chromatin the history layer occupies rather than to identity. Late-replicating, lamina-associated chromatin is the compartment progerin directly disorganises, and it is also where differences in division history accumulate. The cultures were at similar early passages, but their population doublings were not recorded, so this design cannot separate the two.

### Post-irradiation clonal expansion drives methylation away from other cell types, not toward them

Senescent cultures stop dividing, so they cannot show what happens when a damaged cell continues to proliferate. One dataset does: primary fetal fibroblasts were given a single 2 Gy X-ray dose, cloned from single cells, and grown for about twenty population doublings, with non-irradiated clones as controls (Flunkert et al. 2018). The first donor’s control clones match a skin fibroblast line better than any other ENCODE cell type (r = 0.870; Figure 5A), consistent with the panel applying to them. This is the design that should be most favourable to the damage hypothesis.

**Figure 5:**
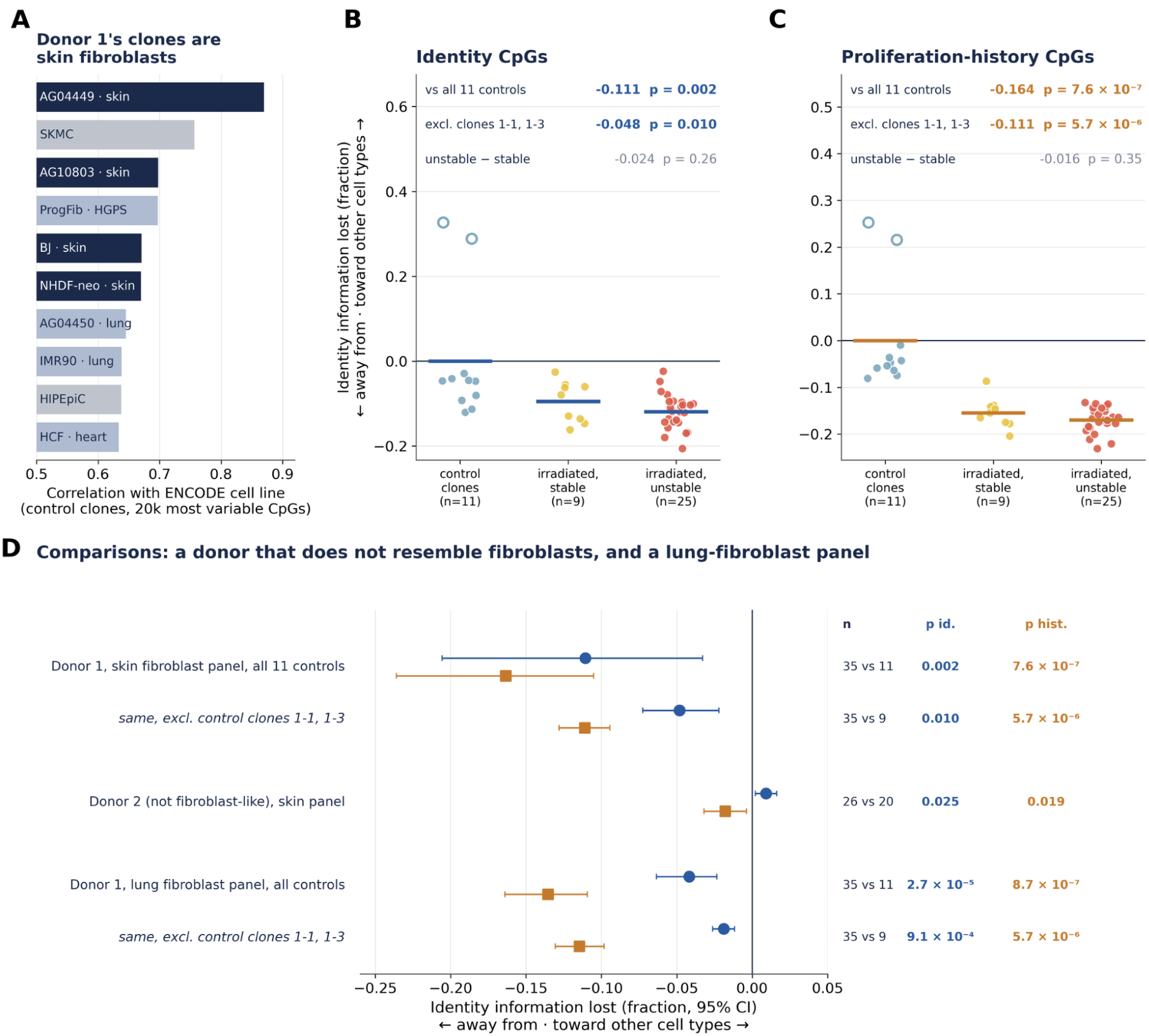
Irradiated clones that kept dividing moved away from other cell types. Single-cell-derived clones of primary fetal fibroblasts, 2 Gy X-ray versus non-irradiated, grown for about 20 population doublings (GSE112812, donor 1). **(A)** The control clones correlate best with skin fibroblast lines (20,000 most variable CpGs). **(B, C)** Identity information lost per clone in the identity and proliferation-history layers, relative to the mean of all 11 control clones. Bars are group means, and open circles are the two outlying control clones. p-values are Mann-Whitney; the unstable-versus-stable comparison is computed in the figure script. **(D)** Controls, with 95% bootstrap intervals. The rows show the same contrast without the two outlier control clones; donor 2 (GSE112873), whose control clones correlate best with renal epithelium; and a lung-fibroblast panel built by the identical procedure, which shares fibroblast-wide CpGs with the skin panel and moves the same way.

Instead, the panel moves in the opposite direction to what the damage hypothesis predicts. Across 35 irradiated and 11 control clones, the identity layer moved away from the other cell types rather than toward them, by 11.1% of its information (p = 0.002), and the history layer moved away by 16.4% (p = 7.6 × 10⁻⁷) (Figure 5B, C). All clones come from one strain, so these p-values describe variation among clones, not among donors. Two control clones are clear outliers at the panel, and excluding them more than halves the identity-layer effect without removing it (−4.8%, p = 0.010). Karyotype instability does not grade the effect: unstable clones differ from stable clones by −2.4% in the identity layer (p = 0.26) and −1.6% in the history layer (p = 0.35). The whole, unsplit panel behaves the same way (Supplementary Figure S2).

Two comparisons bear on whether this divergence is a specific readout of fibroblast identity (Figure 5D). The second donor strain in the same experiment does not resemble a fibroblast in these data. Its control clones correlate best with renal epithelium (r = 0.745) and reach a fibroblast line only at rank 9 (r = 0.641). Its identity-layer score stays close to zero (+0.9%, p = 0.025), nominally significant but a twelfth the size of donor 1’s movement and of the opposite sign. A lung-fibroblast panel built by the identical procedure also moves the same way, but because its comparators exclude every fibroblast, it shares fibroblast-wide CpGs with the skin panel and is not a true negative control. Excluding the two outlier control clones, its identity layer moves 1.9% away from the other cell types (p = 0.0009), against 4.8% for the skin panel, and its history layer moves 11.5% away, against 11.1%. Irradiated clones that keep dividing move away from other cell types at fibroblast-unique CpGs, and we cannot attribute that movement specifically to fibroblast identity.

## Discussion

Across five fibroblast datasets scored on a single quantitative scale, the loss of cell-identity information tracks cumulative cell division. Replicative exhaustion costs 24.3% of the identity information and oncogene activation 5.4%, in cultures that were otherwise identical. A senescence-inducing dose of X-rays costs nothing beyond what baseline-matched background CpGs show, with a 95% interval of −0.7% to −0.2% and a detection limit of 0.4%. Damage-induced senescence by two further triggers does not move the panel consistently. Damage followed by continued division moves the panel further away from other cell types rather than toward them. The simplest reading of these results is that prolonged division, rather than acute DNA damage, is the primary driver of epigenetic identity loss in cultured human fibroblasts. Division is not free of damage, though, and here it is not separated from time in culture: replication stress, telomere attrition and the damage signalling they trigger accumulate over the months that exhaustion takes, so what our contrast separates is acute DNA damage from prolonged division together with the various endogenous damages that accompany it.

Our results do not dispute the phenotypes of the ICE mouse, but they directly question its central interpretation (Yang et al. 2023). In that model, three weeks of non-mutagenic double-strand breaks in an intact animal accelerate epigenetic clocks and tissue aging over the following months, which the authors interpret as repair-driven loss of cell-identity information. Yet epigenetic clocks are not built to measure cell identity, and in a living animal three weeks of breaks followed by months of recovery cannot separate the direct chromatin cost of repair from the cell turnover, compensatory proliferation, unrelated transgene effects, and compositional shifts that tissue injury provokes. Where we isolate the specific CpGs that define a fibroblast and hold the cell type pure, DNA damage, whether acute, chronic, senescence-inducing, or followed through twenty clonal doublings, does not erode identity methylation, whereas cell division erodes a quarter of it. The simplest reading is either that the ICE mouse’s clock acceleration does not reflect loss of cell-type identity at all, or that any identity loss in its tissues was driven by the compensatory proliferation that followed the breaks rather than by repair itself. The direct test, scoring cell-type-unique identity CpGs in ICE tissue, is not yet possible from public data: of 138 sequencing runs in the ICE project, the 15 bisulfite runs are all cultured mouse embryonic fibroblasts, and no genome-wide tissue methylome was deposited. Our data make a clear, falsifiable prediction for that model: at matched break burden, double-strand breaks should erode identity methylation in tissues that divide to repair the injury, and should not erode it in post-mitotic tissues.

The mechanism that makes cell division costly to the methylome is straightforward. Each round of replication requires the maintenance methyltransferase to copy the pattern onto a new strand, and the fidelity of that copy is high but not perfect. Loss accumulates fastest in late-replicating, lamina-associated chromatin, producing the partially methylated domains that mark heavily divided cells and tumours (Zhou et al. 2018), and the same logic underlies mitotic epigenetic clocks (Yang et al. 2016; Teschendorff 2020). Our proliferation-history layer is precisely that phenomenon seen from inside a cell-type panel. It loses large amounts of methylation at UL sites under passaging, gains it at Polycomb-target UH sites, and is shared with heavily divided cell lines by construction. What is new here is the dissociation. In the same cultures and the same passages, the identity layer moves the other way and moves far less, gaining methylation at UL. Division alters two distinct chromatin layers in opposite directions and at very different rates, and an unsplit panel simply reports their sum.

That has a direct methodological consequence for anyone building tissue-unique or cell-type-unique panels, including for our own earlier atlas. Which CpGs enter a panel depends on what is in the comparator set, in a way that is easy to miss. A comparator set containing continuously proliferating lines raises the upper percentile at Polycomb targets and lowers it in partially methylated domains, which silently removes most of the proliferation-history layer and leaves a panel that is almost purely identity. Such a panel works well, but only as an unintended side effect of including transformed lines, and that filtering disappears when the comparators are all primary cells, as ours are. We now think any panel of this kind should be reported with its proliferation-history content made explicit. Scoring how many continuously proliferating cell lines share each CpG’s state is a simple way to do this and requires no external chromatin annotation.

The cross-entropy formulation is what makes the negative result usable. A shift of 0.002 in beta is uninterpretable on its own. A statement that acute X-irradiation costs no identity information beyond matched background, with a 95% interval of −0.7% to −0.2% and a design able to detect a 0.4% loss, is a bounded claim that another experiment can contradict. The subtraction is not optional. Without it, the same X-ray data read as a small, highly significant 1.0% loss that is entirely shared with non-identity CpGs at the same baseline methylation. The subtraction removes that shared change, whatever its cause, so the score measures change beyond it. Any information-loss score built on methylation at extreme sites needs the same control. It also makes panels of different sizes and different beta ranges comparable, which is what lets the identity and proliferation-history layers be read on one axis although they differ twentyfold in size. We are explicit that the measure is a reweighting of the same underlying methylation differences and not a second, independent assay.

Three limitations of the measure itself matter for interpretation. It is a cross-entropy of population beta values, not of molecules, so it reports the average cell and cannot see whether individual cells have become more heterogeneous. Per-cell information loss is a different quantity and needs read-level data. It is signed along one axis, toward the mean comparator cell, so a cell that left its identity in some other direction would score as divergence rather than as loss. And the reference for that generic cell comes from cultured primary comparators, which carry their own passage history, so the axis itself inherits the culture confound. Uncultured, sorted-cell methylomes would remove that last problem and are the obvious next refinement.

All of the evidence above comes from cultured cells, and we deliberately do not extrapolate it to intact tissue. We did score one in vivo contrast, sun-exposed against sun-protected skin from the same people, and keratinocyte identity information does fall there, coherently and across most of the panel (6.9% in epidermis, p < 10⁻⁴; Supplementary Figures S4 and S5). We report it without drawing a conclusion from it. Ultraviolet exposure brings chronic inflammation, epidermal hyperproliferation, immune infiltration and clonal expansion of mutant keratinocyte lineages all at once, and one cohort cannot separate them. The layers do not settle it either. The keratinocyte proliferation-history layer falls by 2.9% (p = 0.17), less than half as much as the identity layer, so this is not the clean dissociation of the culture experiments, but a change in cell composition is not excluded. Settling it would need melanocyte and immune panels we could not build from clean uncultured references, and more than a single cohort.

The mechanism behind the irradiated-clone divergence remains unresolved. Fibroblast-unique CpGs in clones that kept dividing after 2 Gy move away from other cell types, robustly and with large sample-level significance. Two features argue against reading this as a damage effect on fibroblast identity. Karyotype instability, which is the direct readout of residual genome damage in this experiment, does not grade it. Two of eleven control clones are outliers at the panel and account for more than half of the identity-layer effect. A lung-fibroblast panel also responds in the same direction, but it shares fibroblast-wide CpGs with the skin panel, so that comparison cannot show whether the response is specific. Single-cell cloning imposes a severe bottleneck, and the controls are themselves clones, so clonal drift and selection during twenty doublings are plausible alternatives that this dataset cannot exclude. What the experiment does show clearly is the sign: whatever is happening, it is not a collapse toward other cell types.

Progeria is the one ambiguous case in our data, and the ambiguity is worth stating plainly. HGPS fibroblasts move the panel toward other cell types, and persistent DNA damage is one of the best-documented features of the syndrome. But progerin also disorganises the nuclear lamina and remodels lamina-associated domains directly (Köhler et al. 2020), and that is the same chromatin compartment our proliferation-history layer lives in. That layer carries the significant movement here (18.9%, or 21.0% against the four age-matched controls, while the identity layer’s 9.1% and 2.6% are not significant), although the difference between the layers is not itself significant. The cultures were at similar early passages, but without recorded population doublings this dataset cannot separate damage, lamina disorganisation and differences in division history. A comparison at matched population doublings would separate them, and it is a far smaller experiment than the one below.

What would settle the question is a single experiment that is not in the public record. Primary skin fibroblasts would be split into dividing and growth-arrested arms, irradiated with and without ATM inhibition, held at matched population doublings, and profiled both on an array and at single-molecule read level. That design separates damage from division, tests DNA-damage-response dependence directly, and measures both the mean shift and the per-molecule heterogeneity. We looked for existing data that would approximate it. We found no genome-wide methylation dataset with ATM inhibition; the ataxia-telangiectasia methylation data we found come from blood or cerebellum (GSE61133), both confounded by cell composition; and the one published acute-X-ray fibroblast array dataset was never deposited. Settling the question definitively will require performing that experiment directly.

Finally, these results carry a practical implication for studies that measure epigenetic information loss. If identity marks erode mainly as a function of division, then any claim that an intervention preserves or restores epigenetic information has to be compared against a division-matched control, or it will mostly be measuring proliferation. The same applies in reverse to damage models. An intervention that reduces damage usually also changes how much cells divide, and in this system the second effect would dominate the readout. Our data do not say that DNA damage is unimportant in aging. They say that in human fibroblasts, at the specific CpGs that encode what a fibroblast is, acute damage is negligible and prolonged division is the dominant term.

## Methods

### Cell-type-unique CpG panels

We built each panel from GSE40699, the ENCODE/HudsonAlpha Infinium 450K survey of purified human cell lines, which provides 63 arrays covering 61 distinct lines. Replicate arrays of the same line were averaged, giving one value per cell type, which mirrors the percentile-across-types logic of our earlier tissue atlas (Sagy et al. 2025).

A CpG entered a panel as uniquely low (UL) if the target’s mean beta sat at least 0.1 absolute beta units below the 5th percentile of the comparator cell types, and as uniquely high (UH) if it sat at least 0.1 above the 95th percentile. The 0.1 magnitude gate means the site lies outside the range spanned by nearly all comparators, not merely beyond a percentile cut by an arbitrarily small margin.

Comparators were the 20 primary non-fibroblast GSE40699 cell types: AoSMC, HAEpiC, HCM, HCPEpiC, HEEpiC, HIPEpiC, HMEC, HNPCEpiC, HPAEpiC, HRCEpiC, HRE, HRPEpiC, HUVEC, Hepatocytes, NH-A, NHBE, PrEC, RPTEC, SAEC and SKMC. Continuously proliferating lines were deliberately excluded from the comparator set: they hypermethylate Polycomb-target CpG islands, which raises the 95th percentile at the exact sites that define UH candidates. Fibroblast lines were excluded from the comparators of fibroblast panels, so that the panel measures fibroblast identity rather than skin-fibroblast-versus-other-fibroblast.

The skin fibroblast panel used BJ (foreskin), AG04449 (fetal thigh), AG09309 (toe), AG10803 (abdomen) and NHDF-neo (neonatal dermis) as the target. A lung fibroblast panel (IMR90, AG04450) was built identically. Its comparators also exclude all fibroblasts, so it shares fibroblast-wide CpGs with the skin panel and serves as a comparison rather than a negative control. The keratinocyte panel was built from untreated GSE197723 keratinocytes, since GSE40699 contains none, against 21 cell types: the 19 primary non-endothelial types, fibroblasts, and HUVEC. Fibroblasts and HUVEC were profiled in both GSE40699 and GSE197723, and each entered as one centroid averaged over the two batches, so that every comparator cell type counts once. The target also had to clear the same-batch GSE197723 fibroblasts and HUVEC by the same gate. The endothelial panel used ENCODE HUVEC against 21 cell types (the 19 primary non-endothelial types, the two-batch fibroblast centroid and keratinocytes) and had to replicate in GSE197723 HUVEC.

### Identity and proliferation-history layers

For each panel CpG we computed `cancer_side_frac`, the fraction of 30 continuously proliferating GSE40699 lines (A549, BE2_C, CMK, Caco-2, ECC-1, HCT-116, HeLa-S3, HepG2, HL-60, Jurkat, K562, LNCaP, MCF-7, NB4, PANC-1, PFSK-1, SK-N-MC, SK-N-SH, SK-N-SH_RA, T-47D, U87, ovcar-3, NT2-D1, HEK293, MCF10A-Er-Src, GM06990, GM12878, GM12891, GM12892, GM19239) that also sit beyond the target’s own threshold. Not all of these are tumour lines; the set includes HEK293, MCF10A and five lymphoblastoid lines. What they have in common, and the only property we use, is that they have divided far more than any primary culture, so sharing a state with them marks a CpG as a plausible record of proliferation. Identity CpGs have `cancer_side_frac` equal to zero; proliferation-history CpGs have it at 0.2 or above; CpGs in between are retained in the panel file but assigned to neither layer. The thresholds were fixed before scoring any perturbation. Independent-batch replication is the fraction of a layer’s CpGs at which the GSE197723 fibroblast mean also meets the panel rule, clearing the ENCODE comparators’ 5th (UL) or 95th (UH) percentile by the same 0.1 gate. Because it compares the EPIC and 450K platforms, the rate expected by chance is the same test applied to every array CpG outside the panel (0.8% in each direction). Each layer is compared with chance by a binomial test, and identity with history layers by Fisher’s exact test. Both treat CpGs as independent, which neighbouring CpGs are not, so these p-values are approximate.

### Sensitivity of the layer split

The proliferating lines are the only place transformed cells enter this analysis, so we tested whether the split depends on how they are used (`step8_layer_split_sensitivity.py`, `results/panels/layer_split_sensitivity.csv`). Every subset below is scored identically, as the mean shift toward the other cell types beyond baseline-matched background across the 24 matched GSE197723 quadruplets, with UL and UH pooled by CpG count.

Using the same 30 lines among the comparators instead of as a label, so that proliferation CpGs never enter the panel, gives 1,154 UL and 25 UH CpGs and reproduces the identity-layer result: replicative exhaustion +0.082, RAS +0.018 and X-ray −0.002, against +0.108, +0.023 and −0.002 for the identity layer. 348 of its CpGs are identity CpGs. We did not adopt it because it leaves almost no UH CpGs, and so discards the Polycomb half of the measurement.

Three rules that use no proliferating-line data fail to isolate the identity layer. Selecting the CpGs furthest outside the comparator range (top 5%, 458 CpGs) leaves the panel at −0.088 under replicative exhaustion; selecting CpGs where the comparators themselves cluster tightly (IQR bottom 20%, 1,830 CpGs) leaves it at −0.061; and requiring the unique call to reproduce in an independent batch of fibroblasts, by the replication rule above (2,872 CpGs), leaves it at −0.024. All three still move away from the other cell types under passaging, where the identity layer moves toward them. The last recovers 67% of the identity layer but at 8.5% precision, because the proliferation-history layer is fourteen times larger at UL. Only the conjunction of the last two moves toward the other cell types, by +0.019 against +0.108 for the identity layer (595 CpGs, 12% precision), and we treat that as a post-hoc combination rather than a principle.

### Processing of GSE40699

GSE40699 is deposited as raw IDATs with no processed matrix, so we processed it with `methylprep` at its defaults (infer channel switch, quality mask, NOOB). The sample sheet was derived from the series matrix rather than written by hand, emitting one row per Grn IDAT across all supplementary-file fields, which matters because one sample (HCT-116) was arrayed twice. `methylprep` 1.7.1 calls `DataFrame.append`, removed in pandas 2.0, so this step runs in its own pinned environment; no other step depends on it.

### Baseline-matched background

Panel CpGs sit at extreme beta by construction, and extreme beta regresses on its own. For every panel CpG we therefore drew 20 background CpGs from outside the CpGs being scored (the panel layer and direction in question), matched on control-group mean beta in bins of 0.02, with duplicates kept so that the background inherits the panel’s baseline distribution exactly. CpGs with a missing value in any sample of a contrast were dropped before the draw. In the skin cohort the background was drawn once per skin layer against all of its sun-protected samples, so that the age groups are subsets of the same scores. Draws are seeded, so a rerun reproduces every table. Because the pool excludes only the CpGs being scored, CpGs from other layers of the same panel can enter it. Excluding every CpG of all four cell-type panels instead changes no fraction of identity information lost by more than 0.8 percentage points; in GSE197723 the identity layer’s losses become 24.0% under exhaustion, 5.3% under RAS and −0.4% under X-rays (95% CI −0.7 to −0.2, p = 0.018) (step9_sensitivity_checks.py). The excess score for a sample is its mean panel beta minus its mean background beta, signed by direction (`low` negated) so that positive means divergence away from other cell types; this is `excess_shift` in the result tables. The text and figures report its negative, the shift toward the other cell types, so that positive means identity loss on both of the paper’s scales.

### Cross-entropy information loss

For each CpG we computed two reference probabilities: `q_T`, the target cell type’s mean beta, and `q_O`, the mean of the comparator primary types. Treating a CpG as Bernoulli, the extra code length of describing a sample under the identity model rather than the generic one is

`dH_i = H(b_i, q_T) − H(b_i, q_O) = const_i − b_i · (logit q_T − logit q_O),`

which is linear in beta. The constant cancels in any contrast, leaving

information lost (nats) = `Σ_i (b_i^trt − b_i^ctrl) · dlogit_i, with dlogit_i = logit q_O − logit q_T.`

Moving a CpG the whole way from `q_T` to `q_O` costs its Jeffreys term `J_i = (q_O − q_T)(logit q_O − logit q_T) ≥ 0`, so dividing by `Σ_i J_i` gives the fraction of the panel’s identity information that was lost. Zero is unchanged, one means every CpG moved the whole way to the generic cell’s value (before background subtraction), and negative is divergence away from it. Because `dlogit` carries the sign, UL and UH are two halves of one quantity rather than two directions needing a convention, and we combine them weighted by their Jeffreys mass. The identity layer’s mass is 98% UL, so its combined value is in effect a UL measure; the UL and UH halves are reported separately in `information_loss_summary.csv`. Panel directions with fewer than five CpGs are not scored, which excludes only the four endothelial identity UH CpGs.

Probabilities are clipped at 0.02 before the logit, and the whole calculation is repeated at 0.01 to report sensitivity to that choice. Bits per CpG are the fraction times total Jeffreys mass divided by the number of CpGs and by ln 2. Because `loss_i / J_i = Δb_i / (q_O − q_T)`, the fraction is the Jeffreys-weighted mean, over CpGs, of the fraction of the distance from `q_T` to `q_O` that each CpG moved (Figure 3A).

The cross-entropy is corrected for the matched background in the same way as the excess score. For each panel CpG we take the mean methylation change of its 20 background CpGs and weight it by that panel CpG’s own `dlogit_i`. The result is the loss the panel would have shown had every CpG moved like its own baseline-matched background CpGs. That background term is subtracted from the panel’s own loss, and the difference is the fraction reported throughout. The two are computed on the same complete-case CpGs and the same background draw as the excess score. Both components are also reported for every contrast. The background has to be weighted by the panel’s `dlogit` rather than its own: background CpGs carry almost no identity information, so scoring them with their own weights returns a value near zero however far they move.

### Datasets and contrasts

**GSE197723** (Kabacik et al. 2022), EPIC. 24 matched quadruplets of primary fibroblasts, each an untreated control plus a replicatively exhausted, an X-irradiated and a RAS-expressing culture; scored as within-quadruplet pairs. GEO gives no donor field, so we checked the quadruplets with genotype-like calls at 679 probes whose beta is discrete (within 0.08 of 0, 0.5 or 1) in every sample and varies among the series’ 17 skin donors (`step3_cohorts.py`): the four samples of each quadruplet share a genotype (at most 0.3% discordant calls), and 23 of the 24 appear to come from different donors (1.8 to 3.1% discordant), with quadruplets 229 to 232 and 233 to 236 sharing one; we kept all 24, and dropping either of that donor’s quadruplets changes no conclusion (identity information lost under exhaustion 24.1% or 24.7%, under X-rays −0.4% or −0.5%). Each condition occupies the same two array rows on every chip (controls R01 and R05, exhaustion R02 and R06, X-ray R03 and R07, RAS R04 and R08), so pairing cannot remove a row effect; averaging the two quadruplets on each chip leaves the X-ray result unchanged (−0.5%, signed-rank p = 0.034 over 12 chips). The series also includes a four-point untreated and chronically irradiated fibroblast passaging course with recorded population doublings, alongside a three-point keratinocyte equivalent. Untreated keratinocytes and HUVEC from this series provide the reference and replication sets for the keratinocyte and endothelial panels.

**GSE227160**, EPIC. Two dermal fibroblast donors (DSF2, DSF3), each with a young culture, a middle-passage culture, replicative senescence, doxorubicin-induced senescence and irradiation-induced senescence. One young culture per donor serves as reference, so these arms are descriptive and we report no p-values for them.

**GSE149960** (Köhler et al. 2020), EPIC. Nine Hutchinson-Gilford progeria and six control dermal fibroblast cultures. The controls are two parental lines from the Progeria Research Foundation and four age-matched lines from the Coriell repository, and the source study used all of them at early passage (8 to 9 and 9 to 12 passages); population doublings are not reported.

**GSE112812 and GSE112873** (Flunkert et al. 2018), 450K. Primary fetal fibroblasts from two donors, given a single 2 Gy X-ray dose or left untreated, cloned from single cells and grown approximately 20 population doublings, with karyotype stability recorded per clone. Beta values are in the series supplementary files rather than the series matrix, keyed by sample position in series-matrix column order. Four starting-culture samples are not used by any contrast. Two control clones of donor 1 (GSM3084161, GSM3084174) are outliers at the fibroblast panel, and every contrast is reported with and without them. One sample (GSM3084195) is irradiated by its group and description fields but titled as a control clone and has no stability call, and the series design lists 12 control and 34 irradiated clones, so it is probably a control. We kept the group field; excluding the sample or treating it as a control changes the identity-layer difference by at most 0.5 percentage points.

**GSE51954** (Vandiver et al. 2015), 450K. Sun-exposed and sun-protected epidermis and dermis from the same people, across two age groups.

### Which donor is a fibroblast

The clone experiment describes both strains only as fetal fibroblasts, so we checked their resemblance to reference cell types from the data. For each donor we correlated the mean of its untreated clones against every GSE40699 cell type over the 20,000 most variable CpGs. Donor 1 matches the skin fibroblast line AG04449 at rank 1 (r = 0.870). Donor 2 matches renal epithelium HRE at rank 1 (r = 0.745) and reaches its first fibroblast line at rank 9 (HCF, r = 0.641). We therefore report donor 2 separately rather than as a replication. A correlation of this kind flags a potential mismatch with the reference cell types; it does not establish a strain’s lineage.

### Recovering the skin pairs

GSE51954 samples the same person twice but does not say which samples belong together, and the title indices are mis-assigned in young sun-protected epidermis. We recovered the pairs from genotype. Probes with a trimodal beta distribution across samples were selected as SNP-like, samples were correlated on those probes within each layer and age group, and the assignment was solved exactly with the Hungarian algorithm. Pairs with correlation below 0.9 were dropped, leaving 19 dermal and 18 epidermal pairs out of 20 and 19 candidates.

### Population-doubling adjustment

The chronically irradiated cultures of GSE197723 sit 5 to 10% lower in population doublings than their untreated counterparts, and passaging alone moves the panel. We therefore interpolated the untreated passaging series of the same experiment to each irradiated culture’s doubling level and subtracted it. Both the raw shift, against the untreated culture of the same passage, and the adjusted shift are reported.

### Statistics

Paired designs were tested with the Wilcoxon signed-rank test on within-pair differences, unpaired designs with the Mann-Whitney U test, both two-sided, with the sample or pair as the unit of replication throughout. Contrasts with fewer than five units were not tested and are reported as descriptive. Probe-level tests used the Wilcoxon signed-rank test across pairs per CpG, with Benjamini-Hochberg control of the false discovery rate. Confidence intervals on the fraction of identity information lost are percentile bootstraps of the mean over 10,000 resamples, resampling pairs for paired designs and both groups independently for unpaired ones; they are reported alongside, not instead of, the rank tests, and the two can disagree in the smallest arms. Minimum detectable effects quoted in the Results use the standard normal-approximation formula at two-sided α = 0.05 and 80% power, with the observed per-pair or pooled standard deviation. Both the bootstrap intervals and the minimum detectable effects hold the panel, weights and background draw fixed, so they describe sampling of cultures, not uncertainty in those choices. Densities in Supplementary Figure S5 are Gaussian kernel estimates. With thousands of CpGs, probe-level comparisons reach significance at negligible effect sizes, so we treat panel-versus-background differences below 0.005 as uninformative and grey them in the figures.

## Code and data availability

All six datasets used are public. Five are used directly from their GEO series matrix; GSE40699 requires the one-off IDAT processing described above. The analysis is a numbered pipeline of nine steps that rebuilds every table and figure in this paper from the raw data in about twenty minutes, with seeded background draws so that reruns reproduce the tables exactly. Figures read effect sizes and p-values from the result tables, so a figure cannot disagree with a table; the two exceptions, the unstable-versus-stable clone comparison and the skin carry-over regression, are computed in the figure script from the per-sample tables. Code, intermediate tables and figure-generation scripts are at the project repository https://github.com/TheBarLab/identity-methylation-division-history.

## Acknowledgments

We thank the Bar lab members for comments and suggestions. LLMs were used to edit and polish text and to assist with scripts for data processing, statistical analyses, and figure preparation. The authors determined the study design and analytical approach, reviewed the data and results, and are responsible for the manuscript’s final wording and interpretation. This work was supported by the Israel Science Foundation (grants 654/20 and 632/20 to DZB). The funders had no role in the study design, data collection, analysis, interpretation, manuscript preparation, or decision to submit.

## Supplementary figure legends

**Supplementary Figure S1:**
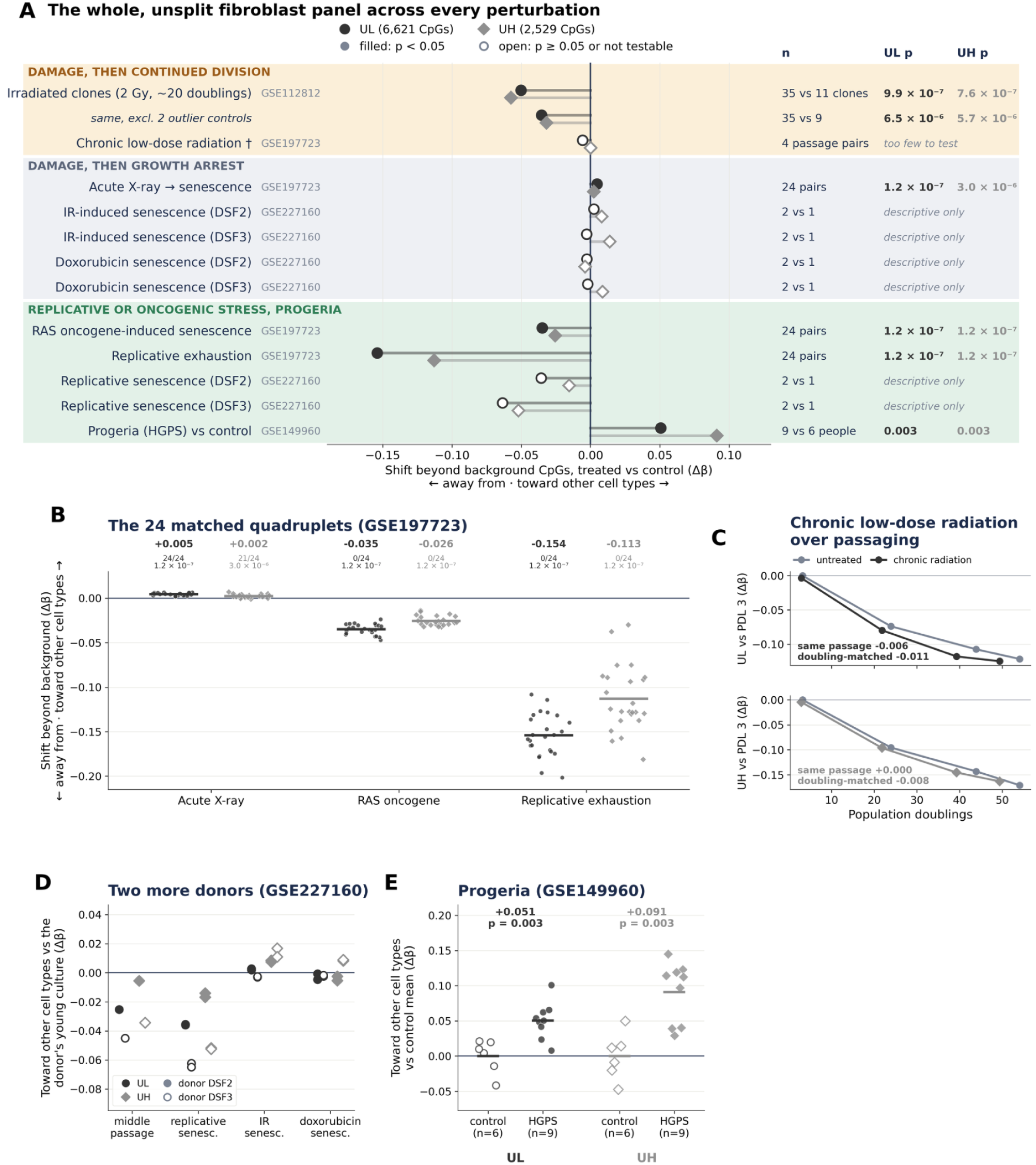
The whole, unsplit fibroblast panel in every contrast. All 9,150 CpGs (6,621 UL, 2,529 UH); shift beyond baseline-matched background in beta, positive toward the other cell types. Most of the panel is proliferation-history CpGs (Figure 2), so these values mostly report division. **(A)** Every contrast, grouped as in Figure 4A, with sample-level p-values. Acute X-ray moves the whole panel slightly but consistently toward the other cell types (+0.005 and +0.002); the identity layer alone does not (Figure 2F). **(B)** The 24 matched quadruplets, as in Figure 2A. **(C)** Chronic low-dose irradiation over passaging. Against untreated cultures of the same passage the shift is −0.006 at UL and 0.000 at UH; interpolating the untreated series to the irradiated cultures’ lower doublings gives −0.011 and −0.008. **(D)** Two further dermal fibroblast donors by senescence trigger, each against its own young culture. **(E)** Progeria versus control dermal fibroblasts.

**Supplementary Figure S2:**
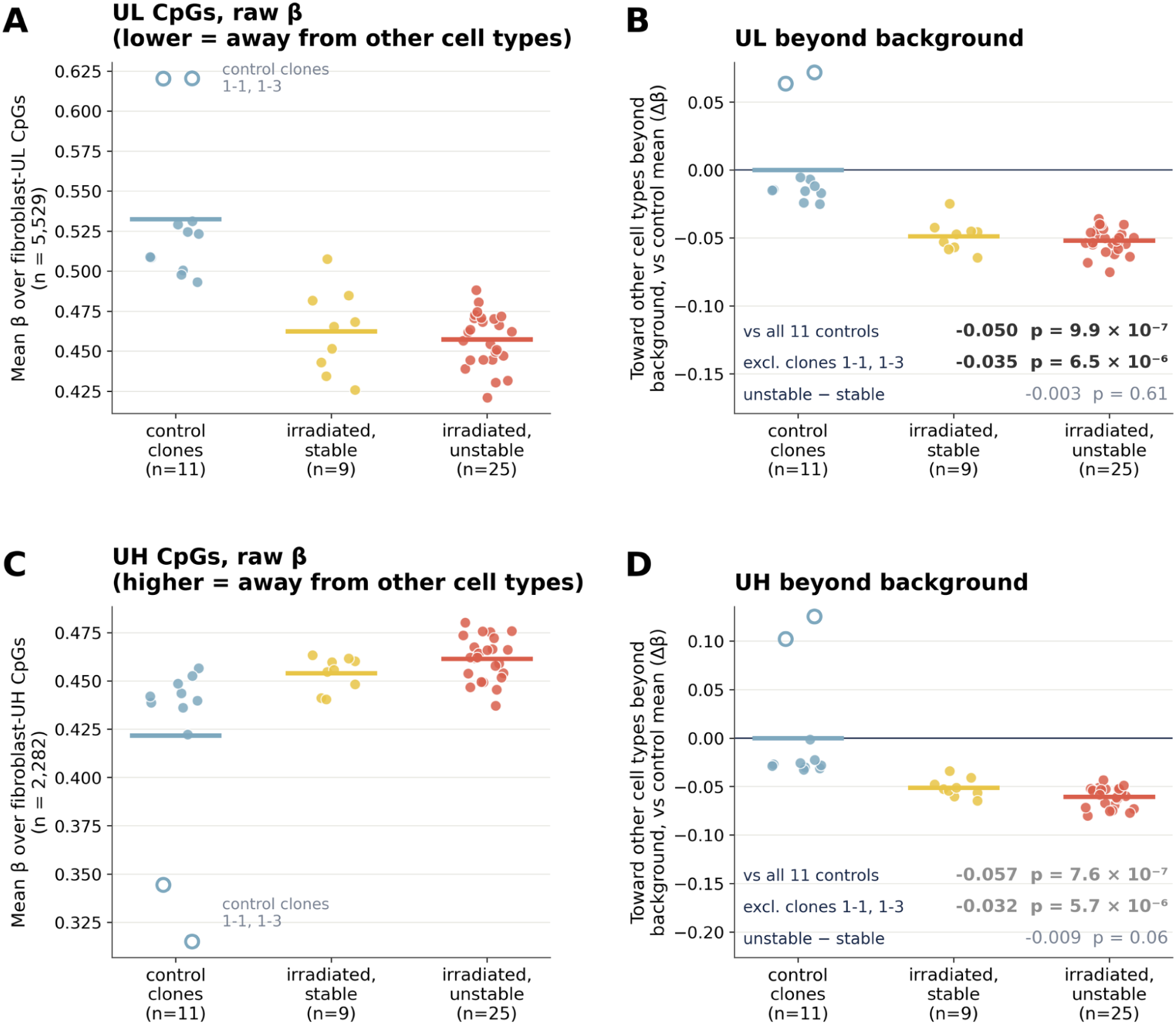
The irradiated clones of Figure 5, clone by clone, on the whole panel. The raw data behind Figure 5 (donor 1, GSE112812), using all 9,150 panel CpGs. **(A, C)** Mean panel beta per clone at UL and UH, by irradiation and karyotype stability. The two outlying control clones (open circles) are outliers in raw methylation, not only after background subtraction. **(B, D)** The same beyond matched background, positive toward the other cell types: −0.050 at UL and −0.057 at UH against all 11 control clones, and −0.035 and −0.032 without the two outliers, with no difference between karyotypically stable and unstable clones (Mann-Whitney; the unstable-versus-stable comparison is computed in the figure script). Bars are group means.

**Supplementary Figure S3:**
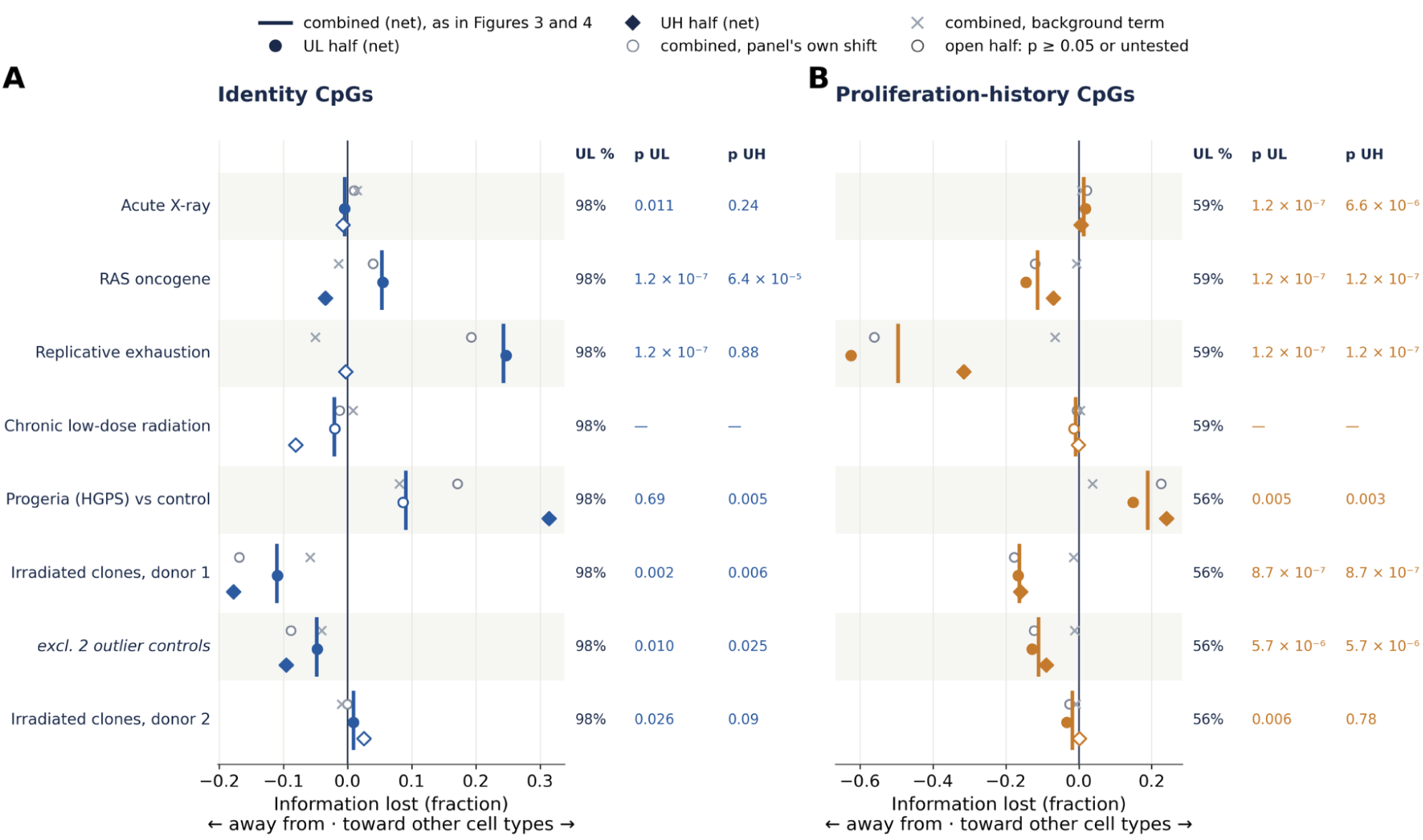
The UL and UH halves of each score, and the background term. For each contrast and layer: the combined score used in Figures 3 and 4 (bar); its UL and UH halves net of background (circle and diamond, filled when p < 0.05); and, for the combined score, the panel’s own shift (open circle) and the background term (×). The columns give the share of the layer’s information carried by UL CpGs and the p-values of the two halves. The identity layer is 98% UL by information, so its combined score is in effect a UL measure; its 10 UH CpGs are shown separately here.

**Supplementary Figure S4:**
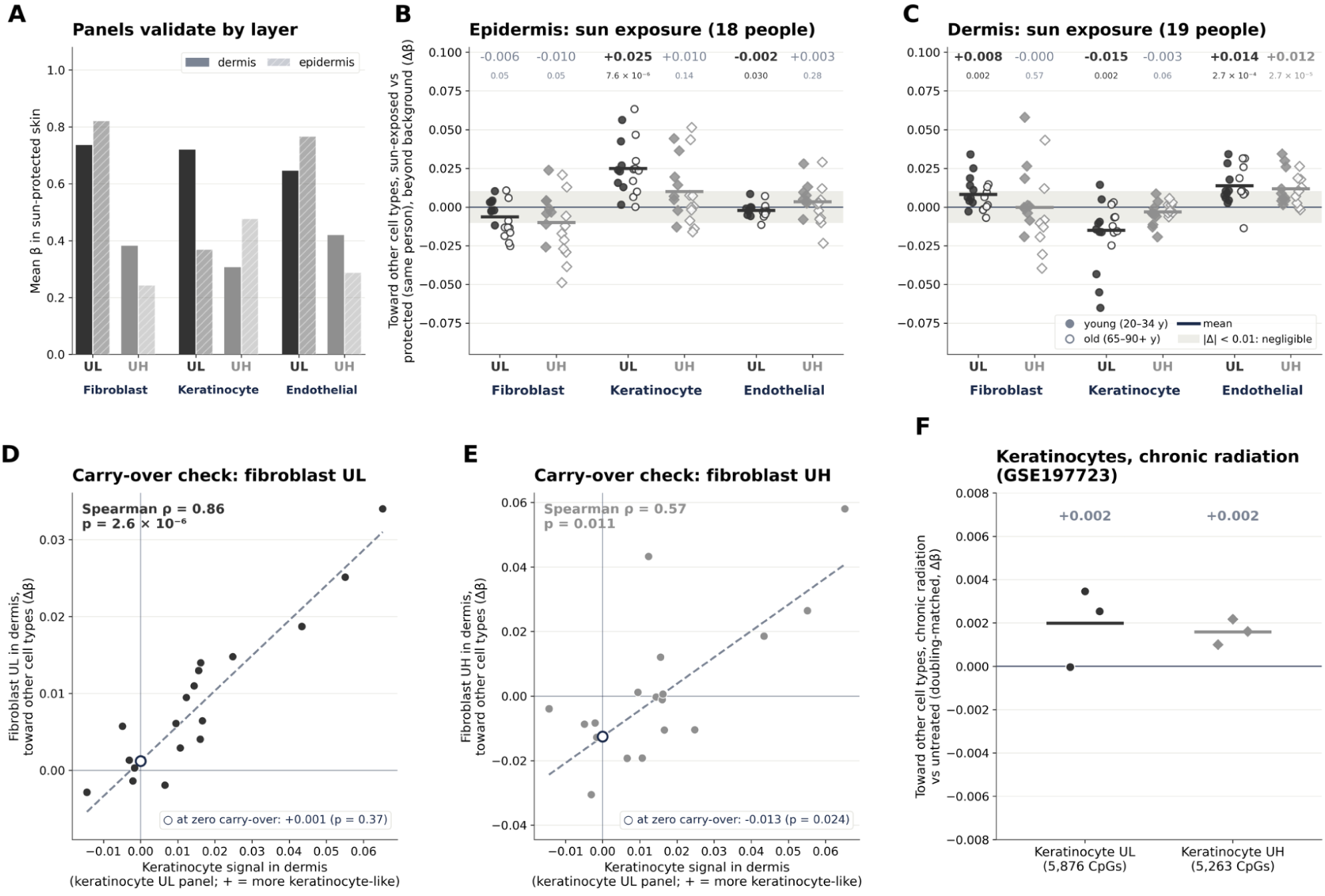
Sun-exposed versus sun-protected skin, paired within person by genotype. Reported for completeness; no conclusion in this paper rests on it (Discussion). Full, unsplit cell-type panels (UL / UH CpGs: fibroblast 6,621 / 2,529; keratinocyte 5,876 / 5,263; endothelial 2,465 / 3,853); shifts beyond background, positive toward the other cell types; numbers above each group are the mean shift and Wilcoxon signed-rank p. **(A)** Each panel validates in the layer that contains its cell type: its UL CpGs are lower and its UH CpGs higher there than in the other layer (dermis for the fibroblast and endothelial panels, epidermis for the keratinocyte panel). **(B)** Epidermis, 18 pairs: the keratinocyte panel moves toward the other cell types (+0.025 at UL, p = 7.6 × 10⁻⁶), while the fibroblast and endothelial panels move by under 0.01. **(C)** Dermis, 19 pairs: the fibroblast panel shifts slightly toward the other cell types (+0.008 at UL, p = 0.002), while the keratinocyte panel moves away from them, indicating more keratinocyte signal in sun-exposed dermis. **(D, E)** Per-person regression of the dermal fibroblast shift on that person’s dermal keratinocyte signal. The UL shift is fully explained by carry-over (ρ = 0.86, fitted shift at zero carry-over +0.001, p = 0.37); the UH shift retains a small residual away from the other cell types (−0.013, p = 0.024). The endothelial panels track the same keratinocyte signal (ρ = 0.52 at UL, 0.69 at UH). The regressions are computed in the figure script. **(F)** Keratinocytes under chronic irradiation, three doubling-matched pairs: too few to test, and all shifts below 0.004. The keratinocyte panel was built from other untreated keratinocytes of the same series.

**Supplementary Figure S5:**
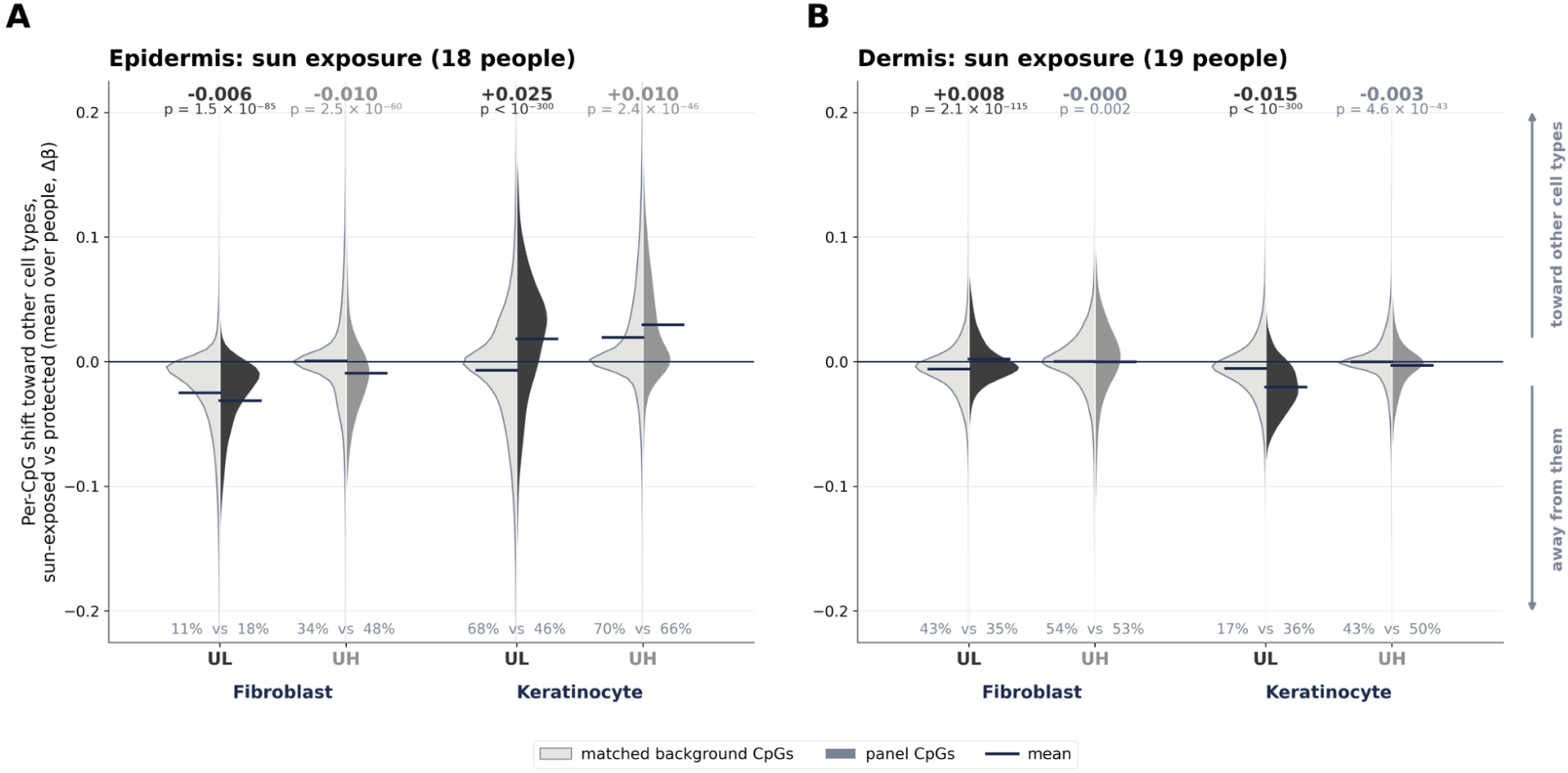
The skin contrast with the CpG as the unit. For each panel CpG, the mean sun-exposed minus sun-protected difference across the genotype-matched pairs of one layer, against the same baseline-matched background the sample-level scores use, signed so that positive is toward the other cell types. Numbers on top: panel mean minus background mean, and Mann-Whitney p; with thousands of CpGs even a negligible difference is significant, so differences under 0.005 are greyed. Numbers at the bottom: share of CpGs moving toward the other cell types, panel against background. **(A)** Epidermis: the keratinocyte UL collapse is coherent. 68% of CpGs move toward the other cell types against 46% of background, and 2,761 CpGs are individually significant at FDR below 0.05 against 934 the other way. **(B)** Dermis: the fibroblast UL shift is a tail rather than a bulk movement (median −0.002, 90th percentile +0.032, 43% of CpGs toward the other cell types against 35% of background). The keratinocyte panel moves away from the other cell types across most of its CpGs (17% toward, against 36% of background), which is the carry-over signal seen at probe resolution.

## References

de Boer J, Andressoo JO, de Wit J, Huijmans J, Beems RB, van Steeg H, Weeda G, van der Horst GTJ, van Leeuwen W, Themmen APN, Meradji M & Hoeijmakers JHJ (2002) Premature aging in mice deficient in DNA repair and transcription. Science 296, 1276–1279.

Di Micco R, Krizhanovsky V, Baker D & d’Adda di Fagagna F (2021) Cellular senescence in ageing: from mechanisms to therapeutic opportunities. Nat Rev Mol Cell Biol 22, 75–95.

d’Adda di Fagagna F (2008) Living on a break: cellular senescence as a DNA-damage response. Nat Rev Cancer 8, 512–522.

Flunkert J, Maierhofer A, Dittrich M, Müller T, Horvath S, Nanda I & Haaf T (2018) Genetic and epigenetic changes in clonal descendants of irradiated human fibroblasts. Exp Cell Res 370, 322–332.

Hoeijmakers JHJ (2009) DNA damage, aging, and cancer. N Engl J Med 361, 1475–1485.

Horvath S (2013) DNA methylation age of human tissues and cell types. Genome Biol. 14, R115.

Kabacik S, Lowe D, Fransen L, Leonard M, Ang S-L, Whiteman C, Corsi S, Cohen H, Felton S, Bali R, Horvath S & Raj K (2022) The relationship between epigenetic age and the hallmarks of aging in human cells. Nat Aging 2, 484–493.

Köhler F, Bormann F, Raddatz G, Gutekunst J, Corless S, Musch T, Lonsdorf AS, Erhardt S, Lyko F & Rodríguez-Paredes M (2020) Epigenetic deregulation of lamina-associated domains in Hutchinson-Gilford progeria syndrome. Genome Med 12, 46.

López-Otín C, Blasco MA, Partridge L, Serrano M & Kroemer G (2023) Hallmarks of aging: An expanding universe. Cell 186, 243–278.

López-Otín C, Blasco MA, Partridge L, Serrano M & Kroemer G (2013) The hallmarks of aging. Cell 153, 1194–1217.

Niedernhofer LJ, Garinis GA, Raams A, Lalai AS, Robinson AR, Appeldoorn E, Odijk H, Oostendorp R, Ahmad A, van Leeuwen W, Theil AF, Vermeulen W, van der Horst GTJ, Meinecke P, Kleijer WJ, Vijg J, Jaspers NGJ & Hoeijmakers JHJ (2006) A new progeroid syndrome reveals that genotoxic stress suppresses the somatotroph axis. Nature 444, 1038–1043.

Oberdoerffer P, Michan S, McVay M, Mostoslavsky R, Vann J, Park S-K, Hartlerode A, Stegmuller J, Hafner A, Loerch P, Wright SM, Mills KD, Bonni A, Yankner BA, Scully R, Prolla TA, Alt FW & Sinclair DA (2008) SIRT1 redistribution on chromatin promotes genomic stability but alters gene expression during aging. Cell 135, 907–918.

Sagy N, Bender O & Bar DZ (2026) Colon-specific epigenetic clocks from minimal features reveal disease-driven aging. Sci. Rep. Available at: https://www.nature.com/articles/s41598-026-64349-3.

Sagy N, Chang C, Gal M & Bar DZ (2025) Epigenetic information loss is a common feature of multiple diseases and aging. GeroScience. Available at: 10.1007/s11357-025-01767-7.

Shtark RB, Sagy N, Korenfeld N, Gal M, Goldstein I & Bar DZ (2026) Reversible and causal epigenetic information loss in liver aging and disease. PNAS Nexus 5, gag109.

Teschendorff AE (2020) A comparison of epigenetic mitotic-like clocks for cancer risk prediction. Genome Med 12, 56.

Vandiver AR, Irizarry RA, Hansen KD, Garza LA, Runarsson A, Li X, Chien AL, Wang TS, Leung SG, Kang S & Feinberg AP (2015) Age and sun exposure-related widespread genomic blocks of hypomethylation in nonmalignant skin. Genome Biol. 16, 80.

Yang J-H, Hayano M, Griffin PT, Amorim JA, Bonkowski MS, Apostolides JK, Salfati EL, Blanchette M, Munding EM, Bhakta M, Chew YC, Guo W, Yang X, Maybury-Lewis S, Tian X, Ross JM, Coppotelli G, Meer MV, Rogers-Hammond R, Vera DL, Lu YR, Pippin JW, Creswell ML, Dou Z, Xu C, Mitchell SJ, Das A, O’Connell BL, Thakur S, Kane AE, Su Q, Mohri Y, Nishimura EK, Schaevitz L, Garg N, Balta A-M, Rego MA, Gregory-Ksander M, Jakobs TC, Zhong L, Wakimoto H, El Andari J, Grimm D, Mostoslavsky R, Wagers AJ, Tsubota K, Bonasera SJ, Palmeira CM, Seidman JG, Seidman CE, Wolf NS, Kreiling JA, Sedivy JM, Murphy GF, Green RE, Garcia BA, Berger SL, Oberdoerffer P, Shankland SJ, Gladyshev VN, Ksander BR, Pfenning AR, Rajman LA & Sinclair DA (2023) Loss of epigenetic information as a cause of mammalian aging. Cell. Available at: 10.1016/j.cell.2022.12.027.

Yang Z, Wong A, Kuh D, Paul DS, Rakyan VK, Leslie RD, Zheng SC, Widschwendter M, Beck S & Teschendorff AE (2016) Correlation of an epigenetic mitotic clock with cancer risk. Genome Biol 17, 205.

Zhou W, Dinh HQ, Ramjan Z, Weisenberger DJ, Nicolet CM, Shen H, Laird PW & Berman BP (2018) DNA methylation loss in late-replicating domains is linked to mitotic cell division. Nat Genet 50, 591–602.

